# Microbial diversity of the glass sponge *Vazella pourtalesii* in response to anthropogenic activities

**DOI:** 10.1101/2020.05.19.102806

**Authors:** Kathrin Busch, Lindsay Beazley, Ellen Kenchington, Frederick Whoriskey, Beate Slaby, Ute Hentschel

## Abstract

Establishment of adequate conservation areas represents a challenging but crucial task in the conservation of genetic diversity and biological variability. Anthropogenic pressures on marine ecosystems and organisms are steadily increasing. Whether and to what extent these pressures influence marine genetic biodiversity is only starting to be revealed. Using 16S rRNA gene amplicon sequencing, we analysed the microbial community structure of 33 individuals of the habitat-forming glass sponge *Vazella pourtalesii*, as well as reference seawater, sediment, and biofilm samples. We assessed how two anthropogenic impacts, i.e. habitat destruction by trawling and artificial substrate provision (moorings made of composite plastic), correspond with *in situ V. pourtalesii* microbiome variability. In addition, we evaluated the role of two bottom fishery closures in preserving sponge-associated microbial diversity on the Scotian Shelf, Canada. Our results illustrate that *V. pourtalesii* sponges collected from pristine sites within fishery closures contained distinct and taxonomically largely novel microbial communities. At the trawled site we recorded significant quantitative differences in distinct microbial phyla, such as a reduction in Nitrospinae in sponges and environmental references. Individuals of *V. pourtalesii* growing on the mooring were significantly enriched in Bacteroidetes, Verrucomicrobia and Cyanobacteria in comparison to sponge individuals growing on the natural seabed. Due to a concomitant enrichment of these taxa in the mooring biofilm, we propose that biofilms on artificial substrates may ‘prime’ sponge-associated microbial communities when small sponges settle on such substrates. These observations likely have relevant management implications when considering the increase of artificial substrates in the marine environment, e.g., marine litter, off-shore wind parks, and petroleum platforms.

## INTRODUCTION

Glass sponges (Hexactinellida) are extraordinary animals with a skeleton made of silicon dioxide and a unique histology which is distinct from all other known sponge classes (Leys *et al.*, 2007). The greatest taxonomic diversity of glass sponges is found between 300 and 600 m depth, with only a few populations occurring in shallow (euphotic) waters (Leys *et al.*, 2007). Hexactinellids are among the most ancient metazoans with an estimated origin of 800 million years ago as determined by molecular clocks (Leys *et al.*, 2007), and therefore represent promising candidates for examining evolutionary ancient biological mechanisms and relationships such as sponge-microbe interactions (Pita *et al.*, 2018). The microbial communities associated with sponges can be very diverse, with more than 63 phyla reported previously (Thomas *et al.*, 2016; Moitinho-Silva *et al.*, 2017). According to current understanding, multicellular organisms can no longer be considered as autonomous entities due to the prevalence of internal symbiotic associations (McFall-Ngai *et al.*, 2013). Instead, the term ‘holobiont’ (syn. ‘metaorganism’ (Bosch and McFall-Ngai, 2011) has been defined to cover the host plus its associated microbiota (Bordenstein and Theis 2015; Rohwer *et al.*, 2002). While the histology and trophic ecology of glass sponges has been a matter of several previous studies (Kahn and Leys, 2017; Kahn *et al.*, 2018), the microbiology of glass sponges still remains largely unexplored (but see Steinert *et al.*, in press, Tian *et al.*, 2016; Savoca *et al.*, 2019).

The glass sponge *Vazella pourtalesii* (Schmidt, 1870) is distributed along the continental margin of eastern North America from the Florida Keys in the southeastern USA to the Scotian Shelf off Nova Scotia, Canada, where it forms pronounced monospecific aggregations with densities reaching up to 4 individuals per m^2^ (**Figure 1**; **Figure 2a**, Kenchington *et al.* unpublished data). In 2013, Fisheries and Oceans Canada (DFO) established two Sponge Conservation Areas (the Sambro Bank and Emerald Basin Sponge Conservation Areas, referred to herein as SCAs) to protect two of the most significant concentrations of *V. pourtalesii* from bottom-fishing activities. However, these closures protect less than 2% of the total area covered by the *V. pourtalesii* sponge grounds (Kenchington *et al.* unpublished data), and bottom fishing activities continue to occur almost immediately adjacent to their borders (DFO, 2017).

**Fig.1.**
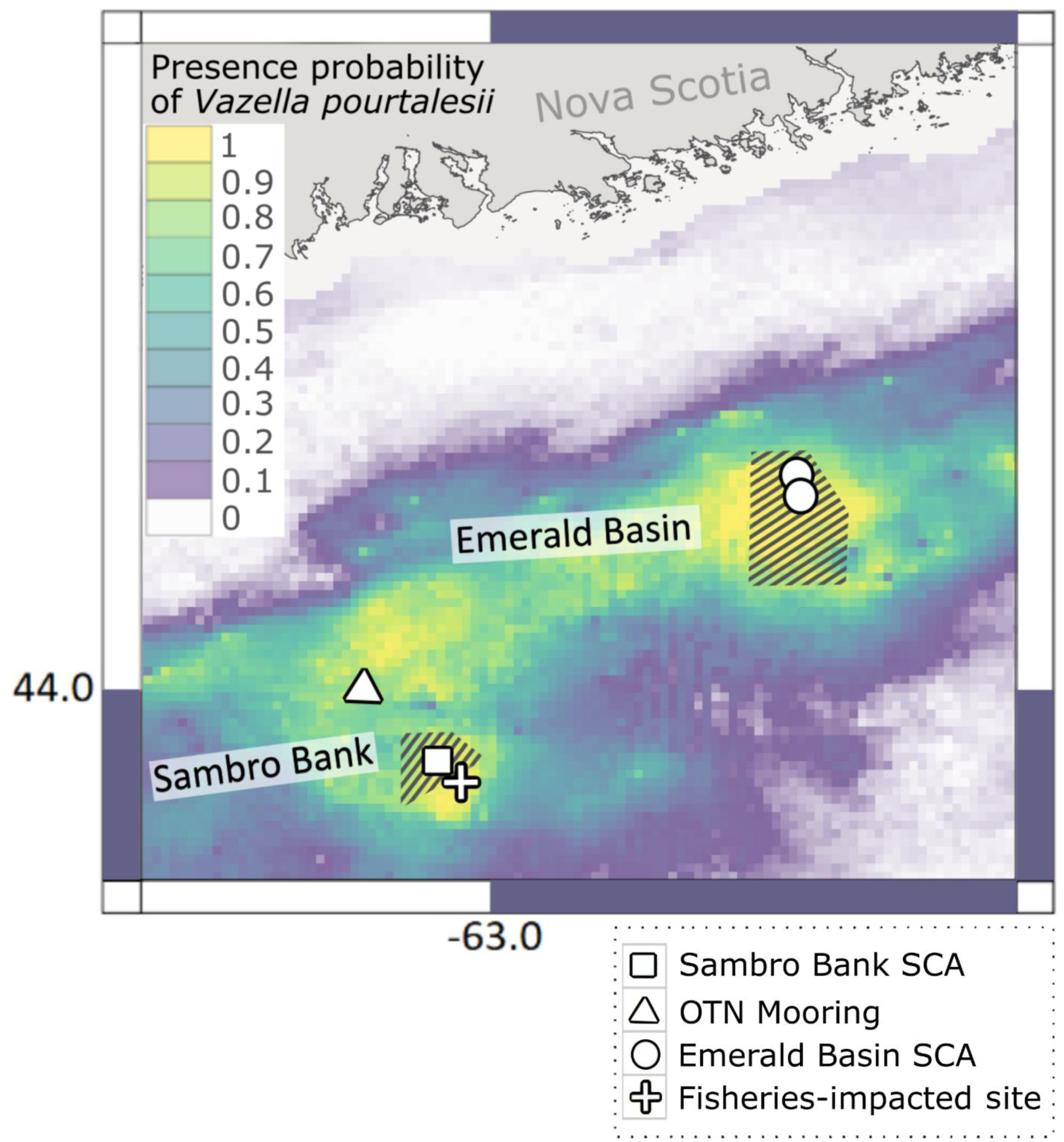
Map of sampling region on the Scotian Shelf, Canada. Colours depict the probability of *V. pourtalesii* occurrence based on the data presented in Beazley *et al.* (2018). Yellow indicates areas where the probability of occurrence is highest. The Sambro Bank and Emerald Basin SCAs are depicted by stripes. Sampling locations of *V. pourtalesii* in the Emerald Basin SCA are indicated by white dots and the white squares at the Sambro Bank SCA. The white triangle depicts the position of the OTN mooring and the white cross represents samples from the fisheries-impacted site

**Fig.2.**
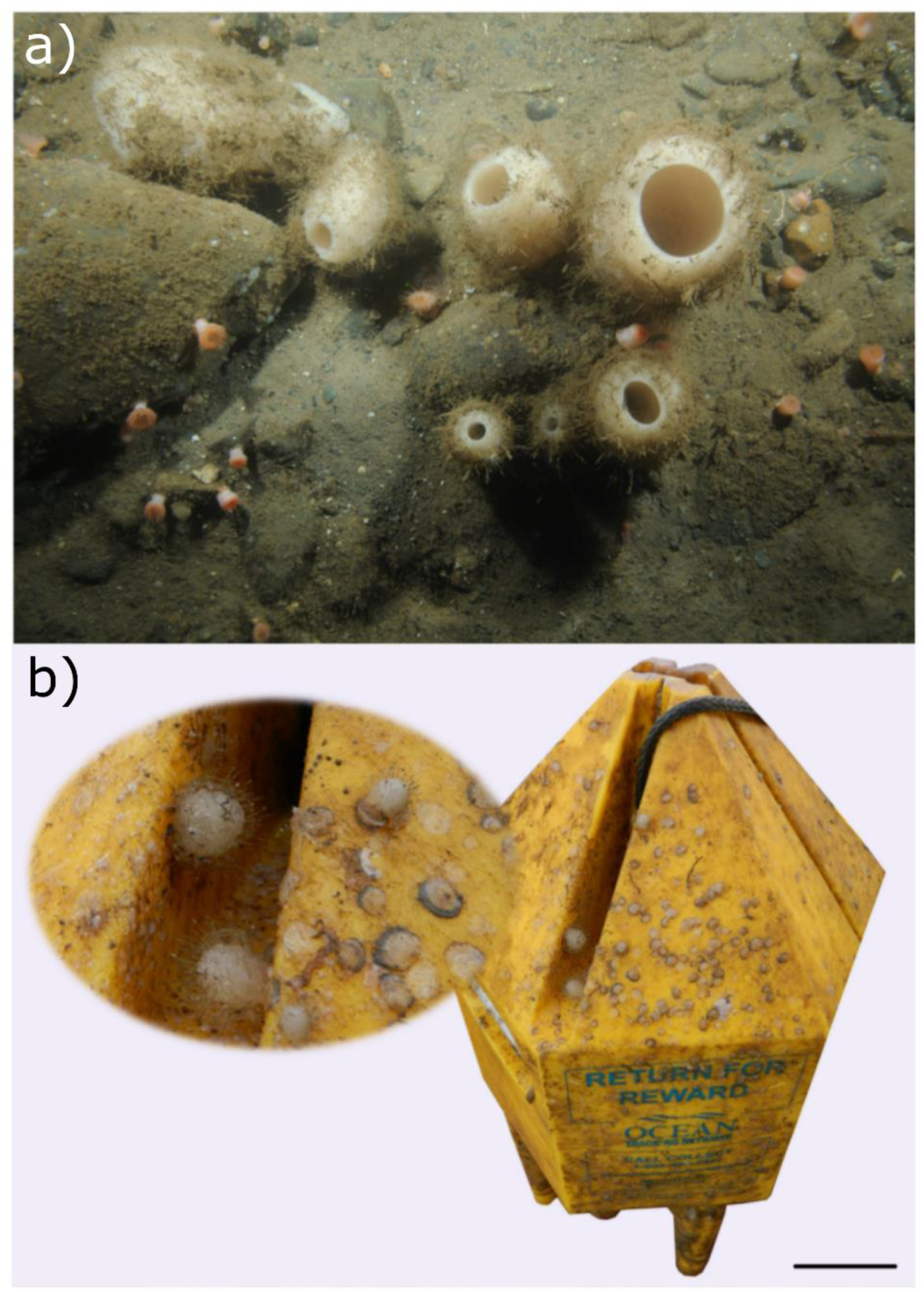
**a)** *In situ* top-view photograph of *V. pourtalesii* aggregation which individuals being up to around 20 cm in height. **b)** Close-up of small *V. pourtalesii* growing on a mooring float of the Ocean Tracking Network (photo credits: DFO). Average size of small *V. pourtalesii* growing on the mooring was approximately 1 cm. Scale bar at right bottom represents 10 cm and applies only to the whole mooring float in Figure 2b

Areas situated in the direct vicinity of fisheries closures represent potentially lucrative areas for fishing, as ‘spillover’ of fish stocks from the protected sites may occur. Bottom trawling is one of the most destructive ways to catch fish (Kelleher, 2005). Impacts include reduced fishing stocks, by-catch of non-target species and destruction of habitat for benthic invertebrates in the trawl path and neighboring vicinity. At trawled sites, the local hydrodynamics may be altered due to a removal of habitat-forming benthic structures (living and non-living). Further, impacts at the population-, community,- and ecosystem-levels have resulted from the removals of unintentionally (bycaught) fished taxa (Ortuño Crespo and Dunn, 2017). Benthic trawling also impacts the microbial community: For example Jackson *et al.* (2001) (and references therein) report a strong increase in microbial cell numbers in the water column associated with overfishing. For glass sponges such as *V. pourtalesii*, the effects of bottom trawling on the variability of microbial community compositions are currently unknown. In terms of sponge physiology, glass sponges were previously observed to arrest pumping in response to high sediment levels in the water column (Tompkins-Macdonald and Leys, 2008), which may have an impact on the oxygenation status inside the sponge tissue and thus on the microbial community.

Several recent studies have explored the response of sponges and their microbiomes (hereafter referred to as ‘holobionts’) to anthropogenic stressors, such as ocean warming, acidification, eutrophication, sedimentation and pollution (reviewed in Pita *et al.*, (2018); Slaby *et al.*, (2019)). The increase in artificial substrates such as plastics in the oceans represents another anthropogenic-induced impact that could potentially adversely affect sponge holobionts. Especially in shelf areas, petroleum platforms, off-shore wind parks and increased loads of litter may represent new settlement opportunities for sponge larvae or gemmules. Artificial substrates have been observed to be exposed to microfouling and biofilm development (Balqadi *et al.*, 2018). It is thus conceivable that biofilms on artificial substrates may influence the sponge holobiont settlement and development on such substrates. The recent discovery that *V. pourtalesii* uses oceanographic moorings (belonging to the Halifax Line of the Ocean Tracking Network (OTN)) as settlement substrate enabled the testing of this hypothesis. The present study aims to describe and compare the microbial community composition of the glass sponge *V. pourtalesii* collected from pristine environments (i.e., within the Sambro Bank and Emerald Basin SCAs), a fisheries-impacted site immediately adjacent to the border of the Sambro Bank SCA, as well as from an artificial substrate (plastic mooring float cover). This study contributes to a deeper understanding of how marine sponge holobionts respond to human pressures.

## METHODS

### Field work

Samples were collected during two oceanographic research missions to the *Vazella* sponge grounds on the Scotian Shelf, off Eastern Canada. The first mission took place in July/August 2016 onboard Canadian Coast Guard Ship (CCGS) *Hudson* (*Hudson2016-019;* Kenchington *et al.*, 2017), where Oceaneering’s Spectrum remotely operated vehicle (ROV) and a mega-box corer were used to collect *V. pourtalesii* from the Emerald Basin SCA (pristine site) between 184 and 206 m depth. The second mission was conducted in September 2017 onboard CCGS *Martha L. Black* (*MLB2017001;* Beazley *et al.*, 2017), where sponge samples were collected during multiple deployments of the ROV ROPOS (Canadian Scientific Submersible Facility, Victoria, Canada) between 153 and 161 m depth at a single station near the centre of the Sambro Bank SCA (pristine site). Furthermore, during this mission, sponge samples were collected outside the closure immediately adjacent to its southeast border at 198 m depth, in an area that was subjected to relatively recent bottom trawling activity (indicated by the presence of deep trawl marks in the general vicinity of the collection site). In total 33 sponges were collected: eight and 13 individuals from the Emerald Basin and Sambro Bank SCAs (pristine sites), and four from the fisheries-impacted site adjacent to the Sambro Bank SCA. Additionally, eight specimens were collected from an Ocean Tracking Network (OTN) mooring located approximately 10 km northwest of the Sambro Bank SCA (for map of sampling locations see **Figure 1**). The mooring (for a picture of the OTN mooring see **Figure 2b**) was anchored ∼ 5 m above the seabed and was deployed for ∼ 13 months (15^th^ of August 2016 - 5^th^ of September 2017) prior to its recovery. Sample metadata were deposited in the Pangaea database.

From each *V. pourtalesii* specimen, pieces of sponge biomass were subsampled and rinsed by transferring them three times through sterile filtered seawater. Excess liquid was then removed and the samples frozen at - 80 °C until DNA extraction. For the mooring samples, complete sponge individuals were frozen at - 80 °C due to their small size.

Seawater samples (2 L) were collected (concomitantly with sponge sampling) at every location in quadruplicates and each filtered onto PVDF filter membranes (Merck Millipore) with a pore size of 0.22 µm and a diameter of 47 mm. Filters were stored at - 80 °C until DNA extraction. Sediments were collected using ROV push corers or box corer deployments at the same locations where sponges and seawater were collected contemporaneously. The top of each core was sliced off, the upper 2 cm of the sediment was collected and stored at - 80 °C until DNA extraction. Biofilms were scraped off the mooring float (where no visible encrusting fauna was present) using cotton swabs (Carl Roth, Karlsruhe, Germany), placed into sterile tubes and frozen at - 80 °C. All sponge, seawater, sediment and biofilm samples covered in this study originated from the same water mass (discussed in Beazley et al., 2018). Throughout this study we use the term ‘environmental references’ for seawater and sediment microbial communities. For assessing the impact of fisheries and artificial substrate provision on *V. pourtalesii* microbial communities, we use Sambro Bank pristine sponges as the baseline for comparison.

### Amplicon Sequencing

DNA extraction was conducted on ∼ 0.25 g of sponge tissue, ∼ 0.25 g of sediment, half a seawater filter or the complete cotton woolen part of a swab using the DNeasy Power Soil Kit (Qiagen, Venlo, The Netherlands). Extracted DNA was quantified using Qubit fluorometer measurements and their quality assessed by a polymerase chain reaction (PCR) with the universal 16S rRNA gene primers 27F+1492R and subsequent gel electrophoresis on 1 % agarose. The V3 and V4 variable regions of the 16S rRNA gene were amplified in a one-step PCR using the primer pair 341F-806R in a dual-barcoding approach (Kozich *et al.*, 2013). The nucleotide sequences of the primers are as follows: 5’-CCTACGGGAGGCAGCAG-3’ (Muyzer *et al.*, 1993) and 5’-GGACTACHVGGGTWTCTAAT-3’ (Caporaso *et al.*, 2011). PCR-products were verified by gel electrophoresis, normalised and pooled. Sequencing was performed on a MiSeq platform (MiSeqFGx, Illumina, San Diego, USA) using v3 chemistry (producing 2 × 300 bp). Demultiplexing after sequencing was based on 0 mismatches in the barcode sequences. Raw reads were archived in the NCBI Sequence Read Archive.

### Bioinformatics and statistics

For analysis of sequencing data, raw sequences were first quality-filtered using BBDUK (BBMAP version 37.75; Bushnell, 2017). Quality trimming was conducted on both ends of the reads (removal of first and last 13 nt) with Q20 and a minimum length of 250 nt. Quality of sequences was evaluated with FastQC (version 0.10.1; Andrews, 2010) and output aggregated with MultiQC (version 0.9; Ewels *et al.*, 2016). The post-filtered sequences were processed with QIIME2 (versions 2018.6 and 2018.8; Bolyen *et al.*, 2018). The DADA2 algorithm (Callahan *et al.*, 2016) was used, which retains Amplicon Sequence Variants (ASVs). Reads were denoised and ‘consensus’ removal of chimera was conducted. One million reads were used to train the error model. Chloroplast and mitochondrial sequences were removed from further analyses. Assignment of taxonomy was conducted using a Naive Bayes classifier (Bokulich *et al.*, 2018) trained on the Silva 132 99 % OTUs 16S database (Quast *et al.*, 2013). We used the term ‘unclassified’ which is defined as ‘no hit’ in this taxonomic classification. This information was used to identify potentially novel microbial diversity in *V. pourtalesii*. Ambiguous classifications (such as ‘unidentified bacteria’) are included in the ‘known sequences’ fraction as those sequence have been recorded and deposited before.

FastTree2 (Price *et al.*, 2010) was used to generate a phylogeny based on ASVs. Subsequently, weighted UniFrac distances (phylogeny-based β-diversity metrices) were calculated (Lozupone *et al.*, 2005). Non-metric multidimensional scaling (NMDS) was performed on weighted UniFrac distances to visually assess sample separation in ordination space. A permutational multivariate analysis of variance (PERMANOVA) was conducted on weighted UniFrac distances to determine whether groups of samples were significantly different from one another. Pairwise tests were performed between all pairs of groups (applied significant level *α* = 0.05). Multivariate analyses were run inside QIIME2 and R.

In addition to phylogeny-based metrics, quantitative measures of community richness were also calculated (e.g. Shannon’s Diversity Index). The Linear Discriminant Analysis (LDA) Effect Size (LEfSe) algorithm (Segata *et al.*, 2011) was applied to determine if microbial phyla differed significantly among groups and rank them according to estimated effect sizes of the significant variations. To accomplish this, first a factorial Kruskal-Wallis sum-rank test was performed on feature tables to detect features (i.e. microbial taxa) with significant differential abundance between groups (applied significant level *α* = 0.05). Wilcoxon rank-sum tests were then run to perform pairwise tests among subgroups (applied significant level *α* = 0.05). Finally, a LDA estimated the effect size of each differentially abundant feature (applied threshold on the logarithmic LDA score for discriminative features = 2.0).

The majority of plots were produced using R statistical software version 3.0.2 (R Development Core Team, 2008) and arranged using Inkscape (version 0.92.3; Harrington and Team, 2005). The *V. pourtalesii* presence probability raster (**Figure 1**) was visualised with QGIS (version 2.18.4; QGIS Development Team, 2017).

## RESULTS

### Microbial diversity of *V. pourtalesii*

In the present study, we assessed the microbial community structure of 33 *Vazella pourtalesii* individuals as well as environmental reference samples by 16S rRNA gene amplicon sequencing. The microbiomes of the *V. pourtalesii* sponges clustered together, and were distinct from seawater and sediment microbiomes (**Figure 3**). Overall, the *V. pourtalesii* microbiomes showed a higher variability of their α-diversity than the environmental reference microbiomes (seawater, biofilm, seawater). Rarefaction curves (α-diversity) revealed further that microbial richness (average number of ASVs per sample) was highest in the biofilm samples collected from the OTN mooring, second highest in seawater, and lowest in *V. pourtalesii* samples (**Online Resource 1: S1**). Also, in terms of diversity indices (Shannon index), alpha diversity was lowest in the *V. pourtalesii* microbiomes. While Proteobacteria were the most dominant phylum in all sample types (sponge: 78.8 % of total; seawater: 47.4 %; sediment: 54.7 %; and biofilm: 56.6 %), Patescibacteria were particularly abundant members of the *V. pourtalesii* microbiome (4.6 %), followed by Bacteroidetes (3.4 %), Spirochaetes (2.9 %), and Planctomycetes (2.8 %), (**Figure 4**). Despite a distinct microbial fingerprint on different taxonomic levels (e.g., at ASV-level, as well as phylum-level), *V. pourtalesii* microbiomes showed a higher variability in terms of their α-diversity than environmental microbiomes. We further observed a higher proportion of previously unrecorded microbial taxa at every hierarchical taxonomic level (microbial phylum down to microbial order level) in the microbiomes of *V. pourtalesii* than in the seawater samples (**Table 1**). At the genus level, 40.2 % of the amplicon sequences of *V. pourtalesii* could not be taxonomically assigned (compared to 16.2 % in seawater). The largest increase of unclassified reads was observed from class to order level (from 0.4 % to 31.6 % respectively). In comparison to environmental microbiomes (seawater, sediment, and biofilm) *V. pourtalesii* microbiomes were particularly enriched in Proteobacteria and Patescibacteria (**Figure 5a**).

**Table 1.** Average fractions of unclassified 16S rRNA gene sequences in *V. pourtalesii* and in seawater at the different hierarchical taxonomic levels indicating a high level of novelty in particularly sponge-associated microbial communities.

| Taxonomic level | Unclassified sequences [%] | Unclassified sequences [%] |
| --- | --- | --- |
|  | in *Vazella pourtalesii* | in Seawater |
| phylum | 0.3 | 0.1 |
| class | 0.4 | 0.3 |
| order | 31.6 | 1.5 |
| family | 36.1 | 10.9 |
| genus | 40.2 | 16.2 |

**Fig.3.**
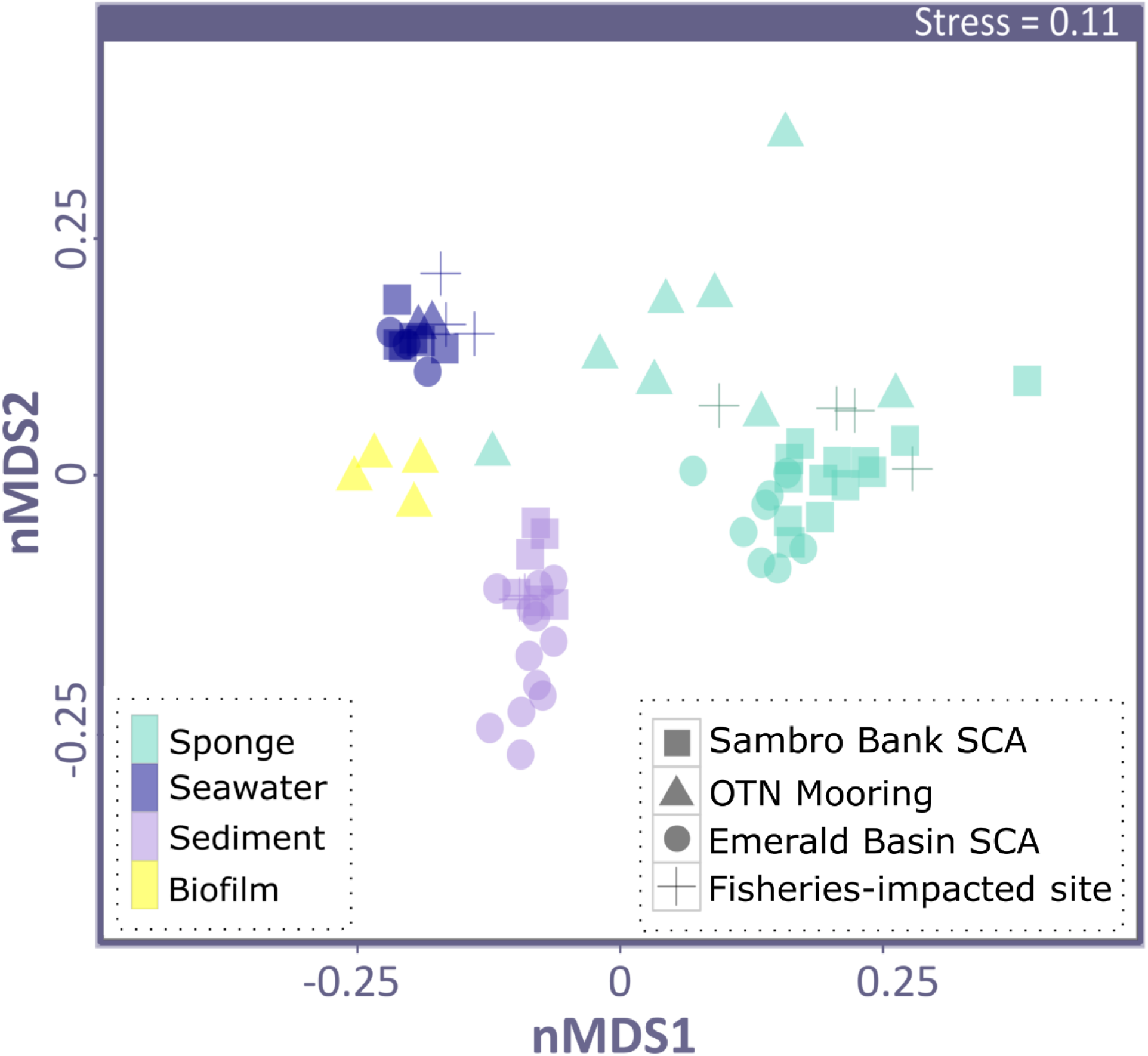
Non-metric Multidimensional Scaling (NMDS) plot on weighted UniFrac distances. Each marker represents a single microbial community. Symbols represent sampling location: squares show samples from Sambro Bank, circles represent samples from Emerald Basin, triangles are mooring samples and crosses are samples from the fisheries impacted site. Colors depict sample type: green marks *V. pourtalesii* samples, blue marks seawater samples, yellow is for mooring biofilm and red represents sediment samples

**Fig.4.**
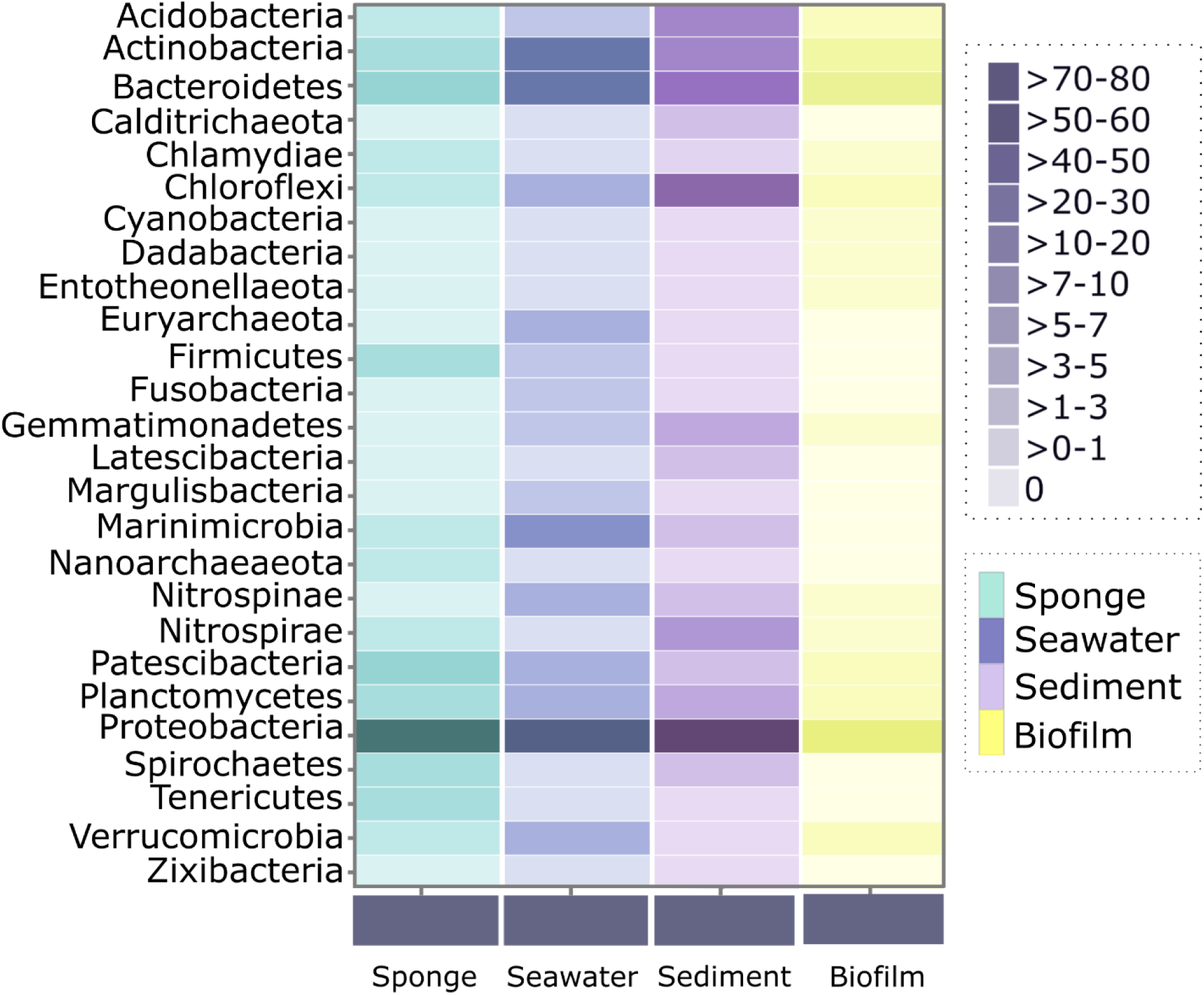
Heatmap showing the relative abundances [%] of microbial phyla per sample type: sponge, seawater, sediment and mooring biofilm

**Fig.5.**
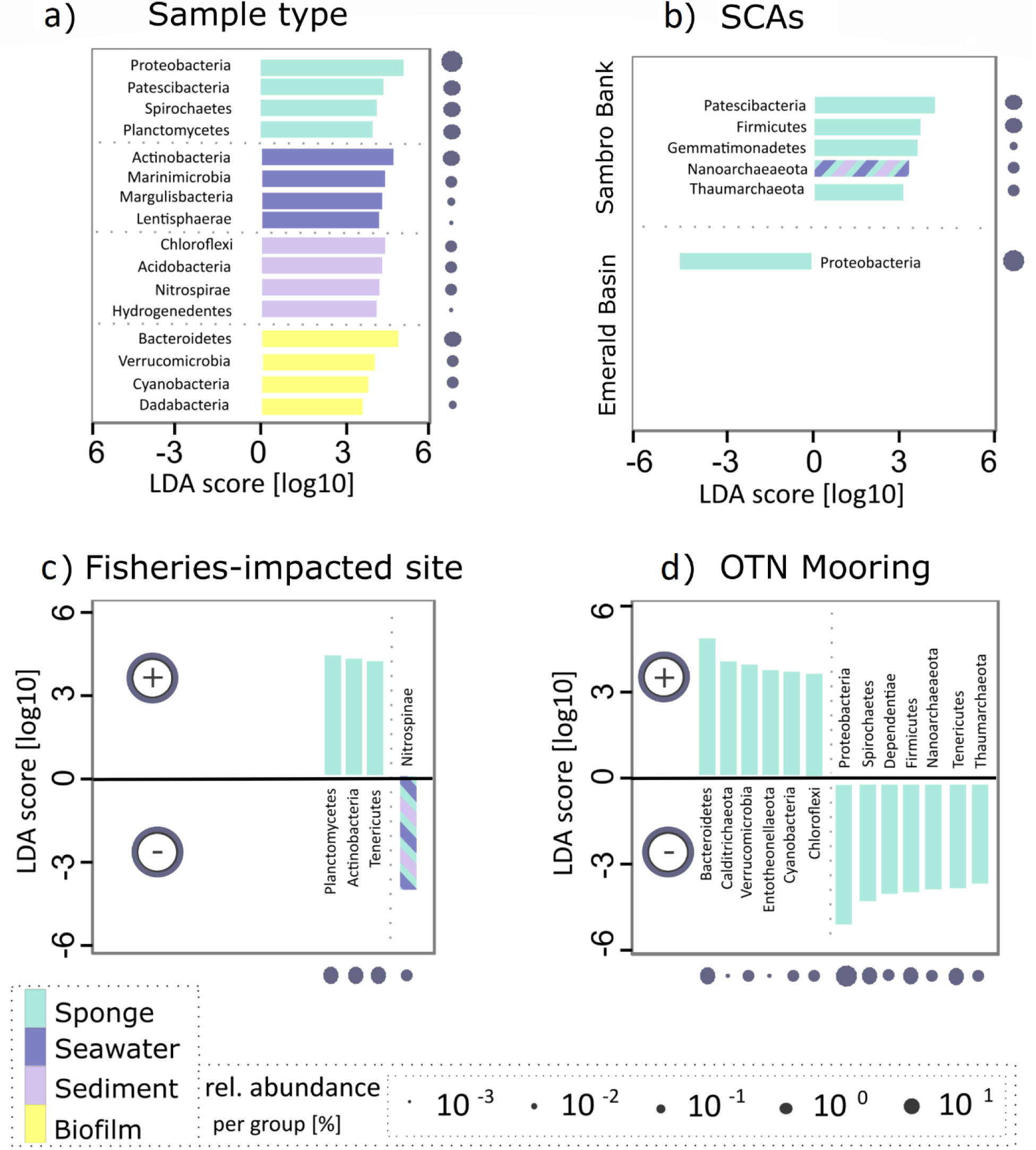
Linear discriminant analysis (LDA) Effect Size (LEfSe) plots. Analysis was performed on microbial phylum level, the rank in the plot is given according to effect size. **a)** Shows data for different sample types (sponge, seawater, sediment and mooring biofilm samples). Here, only the four most significantly different phyla are shown per group. For **b-d)**, where only sponge samples were inspected, all phyla are shown. **b)** Shows differences between sponge conservation areas (SCAs), Sambro Bank and Emerald Basin. c-d) Show microbial taxa increased or decreased upon anthropogenic pressures. **c)** Shows pristine vs. fisheries-impacted sites and **d)** shows growth of sponges in their natural habitat in sponge grounds vs moorings). For all plots, filled bars show microbial taxa with significant effects only in sponge samples; striped bars additionally indicate that these taxa have a significant effect for all three sample types (sponge, seawater and sediment). Dots next to or below plots indicate relative abundance of taxa within the *V. pourtalesii* microbial community. The respective scale is indicated at the lower part of the illustration

### Response of the *V. pourtalesii* microbiome to anthropogenic activities

We then assessed the changes of the sponge microbiome upon human activity, including also implemented protection efforts in form of sponge conservation areas. A significant difference in *V. pourtalesii* microbiomes between the Sambro Bank and Emerald Basin SCAs was observed (**Table 2**), with several microbial phyla significantly enriched in one site over the other (**Figure 5b**). For example, Patescibacteria were significantly enriched in Sambro Bank sponges, while Proteobacteria were significantly enriched in Emerald Basin sponges. We further compared inter-location variability (i.e., *V. pourtalesii* individuals between the Emerald Basin SCA and Sambro Bank SCA) against intra-location variability (i.e., individuals collected from within Emerald Basin SCA only). While the inter-location variability was significant, the intra-location variability was not (PERMANOVA, **Online Resource 1: Table S1**, mean weighted UniFrac distance= 0.29). Further, a smaller mean weighted UniFrac distance was observed between sponges of the same location (Emerald Basin SCA), compared to sponges at different sampling locations (Emerald Basin SCA vs. Sambro Bank SCA). In terms of trawling impact, certain microbial taxa were depleted or enriched in *V. pourtalesii* individuals originating from the trawled site compared to the pristine sites (**Figure 5c**). For example, Nitrospinae were significantly depleted in *V. pourtalesii* individuals originating from the trawled site adjacent to the Sambro Bank SCA (in both sponges as well as environmental references). Nitrospinae are nitrite-oxidising bacteria and have previously been found in deep-sea glass sponges (Tian *et al.*, 2016). With respect to sponge growth on artificial substrates (moorings), their microbial community composition differed significantly from sponges growing in their natural habitat (**Table 2**). Sponges from the artificial mooring substrate were significantly enriched in Bacteroidetes, Verrucomicrobia and Cyanobacteria which were also enriched in the mooring biofilm (**Figure 5d**).

**Table 2.** Pairwise comparisons of beta-diversity are shown for the microbial communities across different sample types, sampling locations, and anthropogenic impacts. Technically, pseudo-F and p-values of pairwise PERMANOVAs on weighted UniFrac matrices are shown.

| group1 | group2 | pseudo-F | p-value |
| --- | --- | --- | --- |
| SambroBankSCA_Sponge | SambroBankSCA_Seawater | 47.65 | <0.01 |
| SambroBankSCA_Sponge | SambroBankSCA_Sediment | 29.76 | <0.01 |
| SambroBankSCA_Seawater | SambroBankSCA_Sediment | 31.35 | <0.01 |
| EmeraldBasinSCA_Sponge | EmeraldBasinSCA_Seawater | 35.62 | 0.01 |
| EmeraldBasinSCA_Sponge | EmeraldBasinSCA_Sediment | 45.33 | <0.01 |
| EmeraldBasinSCA_Seawater | EmeraldBasinSCA_Sediment | 26.87 | <0.01 |
| SambroBankSCA_Sponge | OTNMooring_Sponge | 7.86 | <0.01 |
| Sponge | OTNMooring_Biofilm | 26.37 | <0.01 |
| Seawater | OTNMooring_Biofilm | 18.16 | <0.01 |
| Sediment | OTNMooring_Biofilm | 14.14 | <0.01 |
| SambroBankSCA_Sponge | EmeraldBasinSCA_Sponge | 6.81 | <0.01 |
| SambroBankSCA_Seawater | EmeraldBasinSCA_Seawater | 1.57 | 0.11 |
| SambroBankSCA_Sediment | EmeraldBasinSCA_Sediment | 5.88 | <0.01 |
| SambroBankSCA_Sponge | FisheriesImpactedSite_Sponge | 1.67 | 0.12 |
| SambroBankSCA_Seawater | FisheriesImpactedSite_Seawater | 2.75 | 0.02 |
| SambroBankSCA_Sediment | FisheriesImpactedSite_Sediment | 1.73 | 0.07 |
| FisheriesImpactedSite_Sponge | FisheriesImpactedSite_Seawater | 11.76 | 0.03 |
| FisheriesImpactedSite_Sponge | FisheriesImpactedSite_Sediment | 10.57 | 0.08 |
| FisheriesImpactedSite_Seawater | FisheriesImpactedSite_Sediment | 9.83 | 0.09 |

## DISCUSSION

This study is the first to report on the microbiome composition of the glass sponge *Vazella pourtalesii*, that forms pronounced monospecific aggregations with high densities on the Scotian Shelf, Canada (Fuller, 2011; Beazley *et al.*, 2018) that are not found elsewhere in the world’s oceans. Our results suggest that *V. pourtalesii* contains its own distinct microbial community that is different from seawater and sediment samples. The observed pattern of lower alpha diversity in *V. pourtalesii* compared to environmental reference samples is in line with previous observations showing that sponge-associated microbial communities are usually less complex than those of seawater or sediments (Thomas *et al.*, 2016). We propose that *V. pourtalesii* represents a rich reservoir of novel microbial taxa, even at high taxonomic ranks (e.g. class-level: 31.6 % unclassified). The high abundance of Patescibacteria in *V. pourtalesii* is particularly striking when compared to the microbiomes of other glass sponges (Tian *et al.*, 2016) and also when compared to the environmental reference samples in our study. This microbial phylum is generally assumed to depend on symbiotic animal hosts to cover their basic metabolic requirements (Castelle *et al.*, 2018).

### *V. pourtalesii* microbial community composition varies between the Sambro Bank and Emerald Basin Sponge Conservation Areas

Compared to the conservation efforts towards the protection of animal and plant biodiversity, comparably little is currently known about the needs to protect microbial biodiversity. This is striking as Webster *et al.*, (2018) and others point out very clearly that microorganisms underpin ecosystem health and hence should deserve committed conservation and management endeavors. In our study, we explored the variation in microbial community composition residing in sponges from two SCAs. We observed that the variations in sponge-associated microbiomes were smaller within sites than between sites. We cannot rule out whether a temporal effect may have led to the observed differences between sites, as all Emerald Basin samples were collected in 2016, while all other samples were collected in 2017. However, since no significant differences in seawater microbial communities between both years were observed (**Table 2**), we deduce that temporal effects were not detectable in the water surrounding the sponges. Our results of variable microbial community compositions between Emerald Basin SCA *V. pourtalesiis* and Sambro Bank SCA *V. pourtalesiis* illustrate that fisheries closures are important for the conservation of not only macro-, but also microbial biodiversity.

### Deviations of the *V. pourtalesii* microbial community composition at a trawled site

We observed significantly different relative abundances of distinct microbial phyla from bottom-trawled areas, such as a reduction of Nitrospinae in sponges and reference samples. Their depletion at this fisheries-impacted site is contradictory to our initial expectations. After trawling activities, increased concentrations of nitrite have been recorded in the water column (e.g., Thomas and Kurup, 2004). Pham *et al.* (2019) calculated that removal of sponges by trawling should lead to an increase in nitrite concentrations in the water, due to a lack of uptake by the sponges of this element. We speculate that the observed decrease of Nitrospinae may be triggered by negative feedback loops at the trawled site, such as increased phosphorous or decreased oxygen concentrations. On the other hand, Planctomycetes, Actinobacteria as well as Tenericutes were significantly enriched in *V. pourtalesii* individuals originating from the fisheries-impacted site, but not in the seawater or other samples taken from these sites.

Although we observed significant enrichments or depletions of individual microbial phyla, the overall *V. pourtalesii* microbial community composition did not differ significantly between the pristine and fisheries-impacted site in a permanova analyses based on weighted UniFrac distances. As outcomes of permanova analyses strongly depend on sample sizes, we cannot exclude a technical issue in this regard (as n=4). However, these results are in line with Luter *et al.*, (2012) who observed no shifts in the microbial community of the Great Barrier reef sponge *Ianthella basta* associated with increased sedimentation, when considering the total microbial community rather than individual microbial taxa. In another study on five shallow water sponge species, again no significant effect of increased sediment loads was observed, while minor effects included an increased reliance on phototrophic feeding under high suspended sediment loads (Pineda *et al.*, 2016).

While seawater and sediment microbiomes differed significantly from each other at the pristine sites, microbial community composition was similar for the two sample types at the fisheries-impacted site. Seawater microbial community composition was significantly different at the fisheries-impacted site in comparison to the pristine site, while the sediment microbial community was similar at both sites. From these two observations we conclude that trawling might have an effect on the seawater microbial community in a way that the seawater community becomes more similar to that of sediments. We posit that this may be due to direct and indirect effects of trawling-induced sediment suspension. While a direct translocation of microbial community members may occur from the sediment to the seawater, sediment resuspension may also lead to an increased eutrophication, thus promoting a rise of opportunistic bacterial clades in response to the release of biogeochemical elements (such as e.g., phosphorous) from the sediment into the water column.

### Deviations of the *V. pourtalesii* microbial community composition growing on an artificial substrate

Artificial structures have the potential to act as stepping stones, enabling species to broaden their distributions in the marine environment (Adams *et al.*, 2014). Our results suggest that the biofilms on artificial substrates may ‘prime’ the sponge-associated microbial communities when small sponges settle on such substrates. Our finding of a different microbial community in mooring *V. pourtalesii* individuals compared to pristine ones raises the possibility of these structures changing the microbial diversity of adjacent and newly established benthic populations seeded from such sources. Individuals of *V. pourtalesii* growing on the artificial substrate showed a higher degree of microbiome variation compared to those collected from their natural habitat (**Online Resource 1: S2**). This variation may be due to the fact that sponges growing on the moorings were still small and had presumably settled within a maximum of 13 months following mooring deployment, and thus their microbiomes may not yet have been fully established.

## Conclusions

In the present study two anthropogenic impacts, exposure to trawling activity and growth on plastic moorings, on the *V. pourtalesii* microbiomes were explored. In addition, we evaluated the role of two fisheries closures in preserving sponge-associated microbial diversity on the Scotian Shelf, Canada. We conclude first and foremost, that the Emerald Basin and Sambro Bank Sponge Conservation Areas are ecologically important, as they contain distinct and largely unclassified benthic microbial communities found within sponges or sediments. Variability of the microbial community was comparably high between *V. pourtalesii* individuals, pointing towards the need to minimise sponge loss through anthropogenic pressures (e.g., direct trawling) in order to protect the full range of microbial biodiversity. We further observed significant quantitative differences (in terms of relative abundances) in distinct microbial phyla, such as a reduction in Nitrospinae in sponges and samples from the surrounding environment in areas subjected to bottom trawling. This observed reduction in Nitrospinae is counterintuitive and we suspect that it may be triggered by negative biogeochemical feedback loops. With respect to sponges growing on plastic moorings, significant differences in their microbiome were observed compared to sponges growing on natural substrates. *V. pourtalesii* individuals growing on a mooring surface were significantly enriched in Bacteroidetes, Verrucomicrobia and Cyanobacteria compared to those growing on natural substrates. This could be explained by either the plastic settlement substrate or the younger age of sponges growing on the moorings. Importantly, *V. pourtalesii* is a rich and unique reservoir of microbial biodiversity that deserves conservation effort.

## ACKNOWLEDGEMENTS

This manuscript is written in memory of Hans Tore Rapp, whose research efforts were dedicated to the understanding and preservation of deep-sea sponge grounds in the North Atlantic Ocean. We thank the crews and scientific parties of the two research cruises *Hudson2016-019* and *MLB2017001*. Ulrike Hanz and Furu Mienis are acknowledged for support during field work. We further thank Andrea Hethke, Ina Clefsen, and the CRC1182 Z3 team (Katja Cloppenborg-Schmidt, Malte Rühlemann, John Baines) for assistance with the amplicon pipeline.

## DECLARATIONS

### Ethics approval

Not applicable.

### Consent to participate

Not applicable.

### Consent for publication

Not applicable.

### Availability of data and material

Sample metadata were deposited in the Pangaea database: https://doi.pangaea.de/10.1594/PANGAEA.913907. Raw sequences were archived in the NCBI Sequence Read Archive under BioProject id: PRJNA613976.

### Code availability

Not applicable.

