## Supplementary Material for "Microbial diversity of the glass sponge *Vazella pourtalesii* in response to anthropogenic activities"

related to:

**JOURNAL NAME**

Springer Conservation Genetics

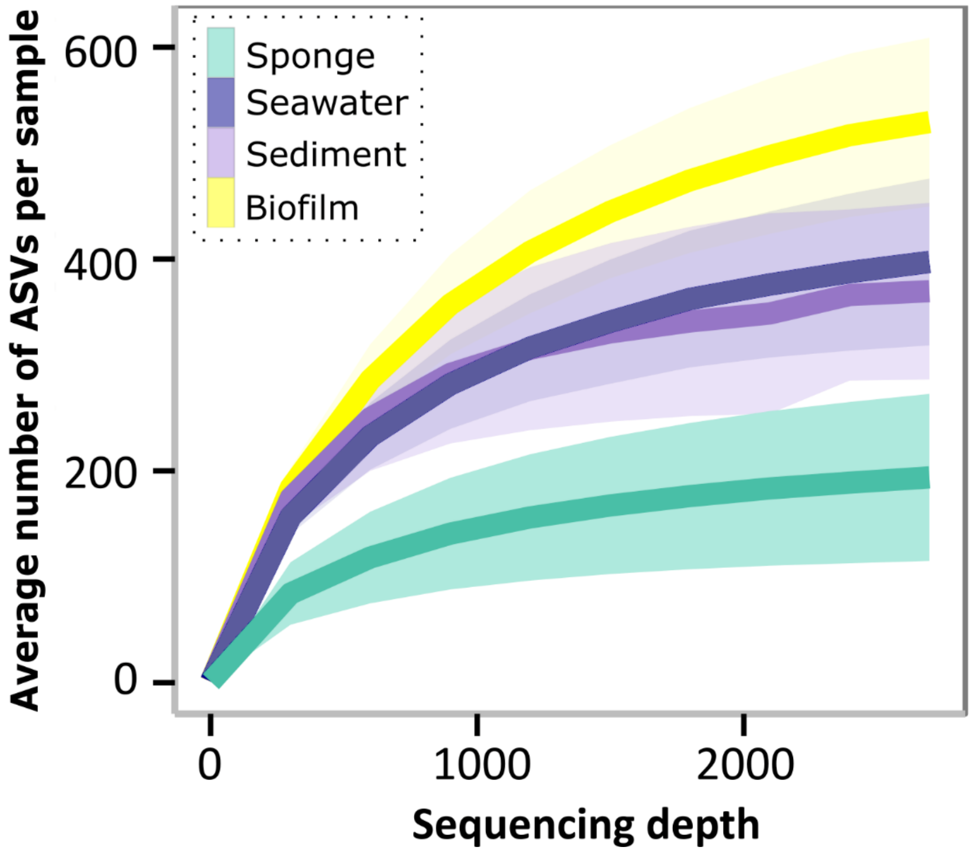


**S1** Alpha-rarefaction curves. Number of observed ASVs per sample is plotted against sequencing depth. Calculated means (solid lines) and standard deviations (colored ranges around lines) of all rarefactions are indicated per sample type (sponge samples - green, sediment samples - purple, seawater samples - blue, biofilm samples - yellow)

**Table S1** Intra-location variation in the microbial community composition of five *Vazella pourtalesii* individuals (B0001 – B0132) from the Emerald Basin SCA is presented. Technically, UniFrac distances between sample pairs and p-values of pairwise PERMANOVAs (999 permutations) are shown

| group1 | group2 | UniFrac distance | *p*-value |
|---|---|---|---|
| EmeraldBasinSCA_Sponge_B0001 | EmeraldBasinSCA_Sponge_B0002 | 0.31 | 0.1 |
| EmeraldBasinSCA_Sponge_B0001 | EmeraldBasinSCA_Sponge_B0003 | 0.23 | 0.1 |
| EmeraldBasinSCA_Sponge_B0001 | EmeraldBasinSCA_Sponge_B0004 | 0.16 | 0.09 |
| EmeraldBasinSCA_Sponge_B0001 | EmeraldBasinSCA_Sponge_B0132 | 0.19 | 0.11 |
| EmeraldBasinSCA_Sponge_B0002 | EmeraldBasinSCA_Sponge_B0003 | 0.33 | 0.33 |
| EmeraldBasinSCA_Sponge_B0002 | EmeraldBasinSCA_Sponge_B0004 | 0.43 | 0.11 |
| EmeraldBasinSCA_Sponge_B0002 | EmeraldBasinSCA_Sponge_B0132 | 0.41 | 0.36 |
| EmeraldBasinSCA_Sponge_B0003 | EmeraldBasinSCA_Sponge_B0004 | 0.31 | 0.18 |
| EmeraldBasinSCA_Sponge_B0003 | EmeraldBasinSCA_Sponge_B0132 | 0.34 | 0.38 |
| EmeraldBasinSCA_Sponge_B0004 | EmeraldBasinSCA_Sponge_B0132 | 0.19 | 0.18 |

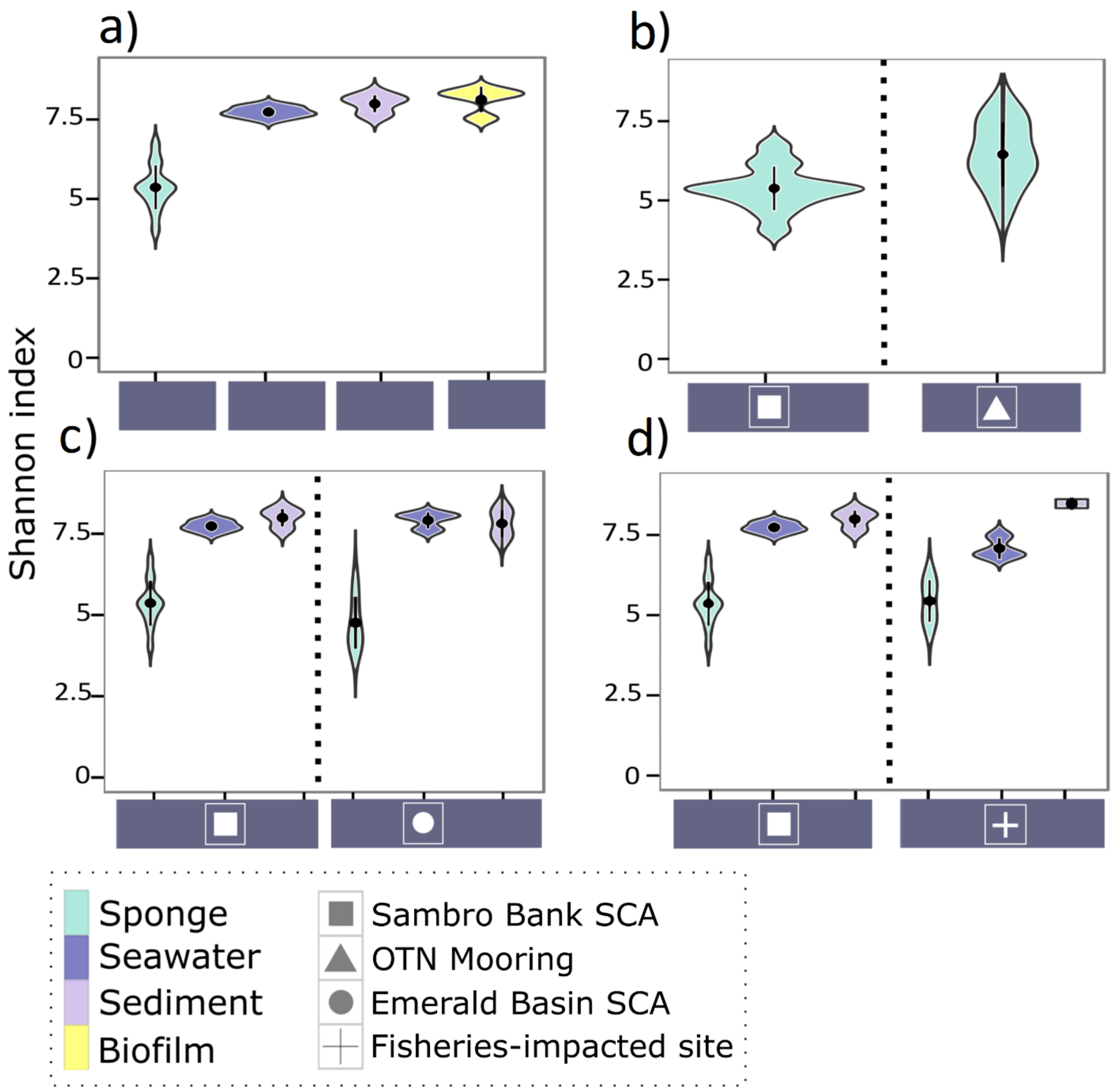


**S2** Microbial community alpha-diversity (Shannon index) of the examined sample types, across locations and habitats. Four graphs show differences between **a)** sample type, **b)** habitat, **c)** sampling area on the Canadian shelf, **d)** fisheries impact. Violin plots show Shannon index of respective samples per group, dots and whiskers inside violins indicate the mean and standard deviation per group
